# Quantifying the efficiency and biases of forest *Saccharomyces* sampling strategies

**DOI:** 10.1101/559724

**Authors:** Primrose J. Boynton, Vienna Kowallik, Doreen Landermann, Eva H. Stukenbrock

## Abstract

*Saccharomyces* yeasts are emerging as model organisms for ecology and evolution, and researchers need environmental *Saccharomyces* isolates to test ecological and evolutionary hypotheses. However, methods for isolating *Saccharomyces* from nature have not been standardized and isolation methods may influence the genotypes and phenotypes of studied strains. We compared the effectiveness and potential biases of an established enrichment culturing method against a newly developed direct plating method for isolating forest floor *Saccharomyces* spp. In a European forest, enrichment culturing was both less successful at isolating *S. paradoxus* per sample collected and less labor intensive per isolated *S. paradoxus* colony than direct isolation. The two methods sampled similar *S. paradoxus* diversity: the number of unique genotypes sampled (*i.e.*, genotypic diversity) per *S. paradoxus* isolate and average growth rates of *S. paradoxus* isolates did not differ between the two methods, and growth rate variances (*i.e.*, phenotypic diversity) only differed in one of three tested environments. However, enrichment culturing did detect rare *S. cerevisiae* in the forest habitat, and also found two *S. paradoxus* isolates with outlier phenotypes. Our results validate the historically common method of using enrichment culturing to isolate representative collections of environmental *Saccharomyces.* We recommend that researchers choose a *Saccharomyces* sampling method based on resources available for sampling and isolate screening. Researchers interested in discovering new *Saccharomyces* phenotypes or rare *Saccharomyces* species from natural environments may also have more success using enrichment culturing. We include step-by-step sampling protocols in the supplemental materials.

## Introduction

Naturally-occurring *Saccharomyces* populations are models for ecology and evolution (Boynton & Greig, 2014). Use of these models has led to exciting discoveries about microbial ecology and evolution; for example, adaptation to climate can lead to speciation (Leducq et al., 2014), domesticated *S. cerevisiae* is more phenotypically diverse than wild *S. paradoxus* (Warringer et al., 2011), and interspecific hybrids can have high fitnesses in stressful environments (Bernardes, Stelkens, & Greig, 2017; Stelkens, Brockhurst, Hurst, Miller, & Greig, 2014). These studies made inferences based on the phenotypes and genotypes of isolates collected from wild and domesticated substrates. And *Saccharomyces* substrates are diverse: wild substrates include tree bark, insect guts, fresh leaves, leaf litter, soil, fruits, and parasitic *Cyttaria* galls, (Glushakova, Ivannikova, Naumova, Chernov, & Naumov, 2007; Kowallik & Greig, 2016; Libkind et al., 2011; Mortimer & Polsinelli, 1999; Sampaio & Goncalves, 2008; Stefanini et al., 2012), and domesticated substrates include wine, beer, bread, kimchi, kombucha, palm wine, and pulque, among many other substrates (Boynton & Greig, 2016; Carbonetto, Ramsayer, Nidelet, Legrand, & Sicard, 2018; Estrada-Godina et al., 2001; Ezeronye & Okerentugba, 2001; Gallone et al., 2016; Greenwalt, Steinkraus, & Ledford, 2000; Jeong, Jung, Lee, Jin, & Jeon, 2013). *Saccharomyces* yeasts are also a single clade in the diverse polyphyletic group of yeasts (single-celled fungi that reproduce by budding or fission) (Kurtzman, Fell, & Boekhout 2011). These diverse yeasts inhabit floral nectar, extreme environments, soils, and insect bodies, among many other habitats (Buzzini, Turchetti, & Yurkov 2018; Chappell & Fukami, 2018; Stefanini, 2018; Yurkov, 2018). One challenge of environmental yeast sampling is to minimize sampling biases so researchers can assure that observed diversity patterns are not artifacts of their chosen sampling strategy.

Enrichment culturing is a reliable and frequently-used method for isolating difficult-to-culture bacteria, archaea, and eukaryotic microbes, including *Saccharomyces*, from natural environments (Korzenkov et al., 2019; Li, Podar, & Morgan-Kiss, 2016; Schlegel & Jannasch, 1967; Sniegowski, Dombrowski, & Fingerman, 2002) (Figure 1A). Microbiologists have been relying on enrichment cultures for over a century (Beijernick, 1961), and have used it to isolate many of the model *Saccharomyces* strains commonly used in laboratory studies (Johnson et al., 2004; Liti et al., 2009; Sniegowski et al., 2002). To isolate a microbe using enrichment culturing, a researcher adds a small amount of natural material to a growth medium designed to be hospitable to the target microbe and inhospitable to other microbes (Liti, Warringer, & Blomberg, 2017; Schlegel & Jannasch, 1967). If the enrichment medium is well-designed, the target microbe is expected to grow in abundance, and after some incubation time, this enrichment culture can be streaked to a solid medium and colonies of the target microbe can be easily isolated. An alternative to enrichment culturing is to spread a microbial substrate directly onto a selective solid medium, with or without dilution, and to pick colonies which morphologically resemble the target microbe (Glushakova et al., 2007; Stefanini et al., 2012) (Figure 1B).

**Figure 1:**
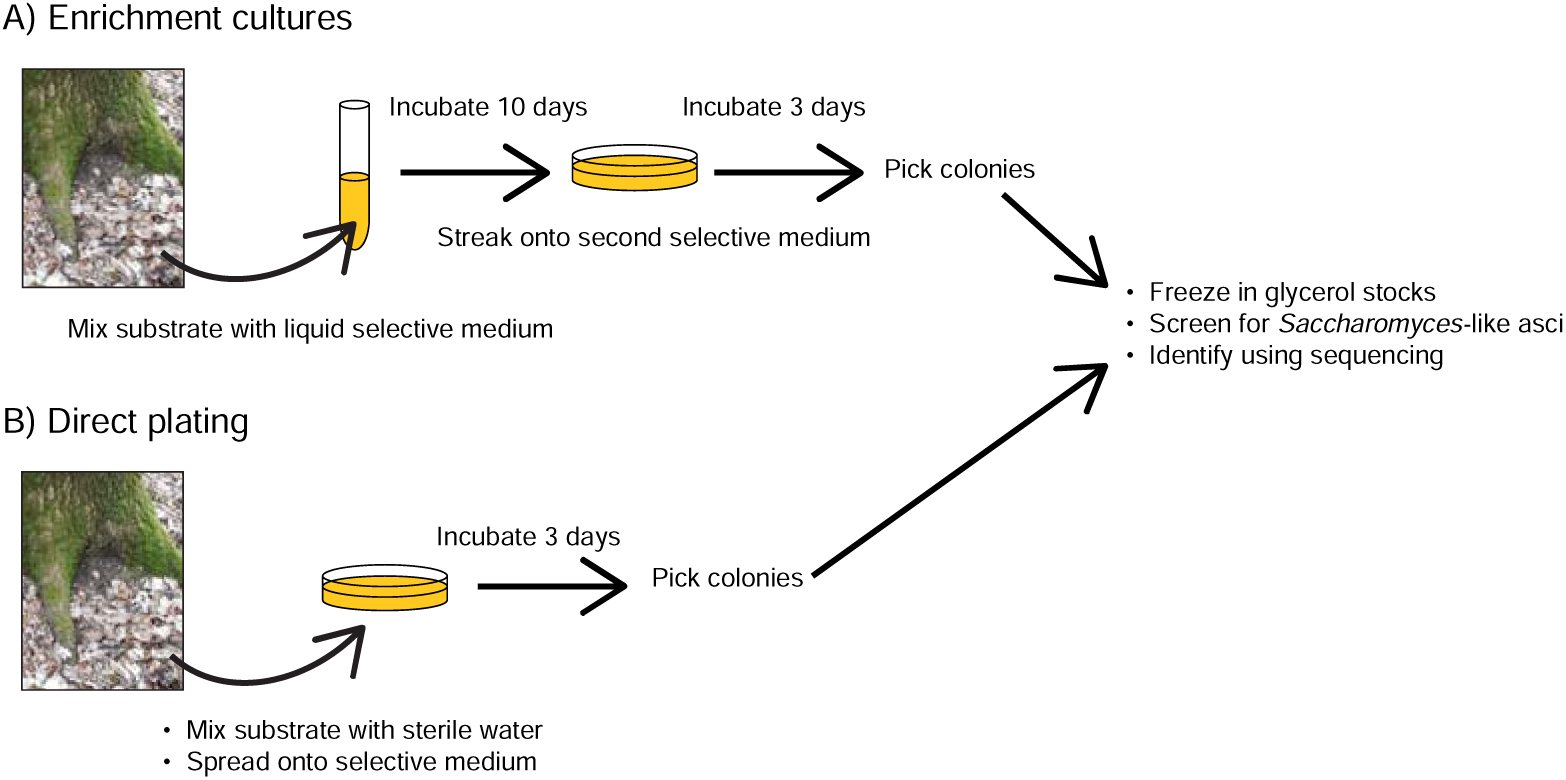
Schematic illustration of sampling strategies used to isolate *Saccharomyces* for this project. A) Enrichment culturing B) Direct plating. Photo: Doreen Landermann.

Because it can be difficult to isolate *Saccharomyces* from natural substrates, many investigations of wild *Saccharomyces* rely on enrichment culturing, usually in high-sugar, acidic media (Charron, Leducq, Bertin, Dube, & Landry, 2014; Robinson, Pinharanda, & Bensasson, 2016; Sniegowski et al., 2002; Sweeney, Kuehne, & Sniegowski, 2004). Comparative studies of *Saccharomyces* genomes have been carried out using collections of *Saccharomyces* strains isolated using various strategies, including both enrichment and direct culturing (Liti et al., 2009; Peter et al., 2018). However, isolation strategy can influence the genotypes and phenotypes of isolated microbes: previous studies have documented higher genotypic diversity among bacteria isolated using direct plating compared to enrichment cultures, and the authors attributed these differences to selection for fast-growing phenotypes during enrichment (Dunbar, White, & Forney, 1997; Oda et al. 2008). We were concerned about the biases that might be introduced during enrichment culturing of *Saccharomyces* yeasts. For example, enrichment culturing might select for individuals with high relative fitness in the enrichment medium. Such potential biases in sampled yeast phenotypes are likely to lead to biases in sampled genotypes because genetic information is responsible for expressed phenotypes. Isolation biases have also been suggested as potential explanations for differences between results of culture-dependent and culture-independent studies of environmental *Saccharomyces* (Alsammar et al., 2018).

This study’s goals were to compare isolation success between enrichment culturing and a direct culturing strategy, and to quantify biases in *Saccharomyces* phenotype and genotype diversity that might be introduced when sampling a forest environment. We tested the assumption that it is easier to sample *Saccharomyces* from forest substrates using enrichment cultures than direct plating. We also investigated potential biases introduced by enrichment culturing by comparing growth rates and sampled genotype diversity between *S. paradoxus* (the wild sister species of the model laboratory yeast *S. cerevisiae*) colonies isolated using enrichment and direct strategies.

Enrichment culturing might decrease or increase sampled *S. paradoxus* diversity compared to direct plating, thereby decreasing or increasing number of genotypes sampled, variance among growth rates, or both. For example, the number of unique genotypes sampled and the variance among growth rates would be low (and average growth rates high) among *S. paradoxus* isolated using enrichment cultures if the enrichment conditions select for the fastest growing *S. paradoxus* genotype present in every sample. Conversely, genotype diversity and variance among growth rates would be high among *S. paradoxus* isolated using enrichment cultures if diversity in the non-*Saccharomyces* microbial communities present on sampled substrates selects for diverse *S. paradoxus* among samples. *S. paradoxus* reproduction during enrichment may also influence sampled genotype diversity: diversity within individual enrichment cultures may be low if a fast-growing genotype makes many asexual copies of itself, or unique genotypes may be produced during enrichment culturing if *S. paradoxus* individuals sexually outcross with one another.

To test these predictions, we compared *Saccharomyces* sampling success and phenotype and genotype diversity among soil and leaf litter samples from a well-studied northern German forest (Kowallik & Greig, 2016; Kowallik, Miller, & Greig, 2015). A previous study showed that *S. paradoxus* is readily isolated using enrichment cultures from oak leaf litter in this forest (Kowallik & Greig, 2016). We were also previously able to isolate *S. paradoxus* directly from these forest substrates without enrichment (Kowallik, 2015). For the current study, we adapted a frequently used published enrichment method, which includes an enrichment step and two selective media, to design a direct plating method which included no enrichment steps and only one selective medium (Figure 1) (Kowallik & Greig, 2016; Sniegowski et al., 2002). We aimed to remove as many potentially bias-inducing steps for the direct plating method, while still being able to isolate *Saccharomyces* spp., to understand whether these commonly-used selective steps bias environmental *Saccharomyces* sampling.

## Methods

### Field sampling and yeast isolation

All isolates were sampled from a mixed hardwood and conifer forest in Nehmten, Schleswig-Holstein, northern Germany (Nehmtener Forst). We sampled approximately seven compressed ml total of each of leaf litter and soil from close to the bases of ten oak trees at four sampling dates (Table 1), although not all trees were sampled at every date. Trees were between 12 and 744 m from one another. At each date, samples were collected from leaf litter and the top organic layer of soil within one meter of the base of each tree. Paired leaf litter and soil samples were collected on the north, south, east, and west side of each tree at all collection dates except 7 April, when samples were collected at an arbitrary two of the four cardinal directions.

**Table 1:** Number of sampling points from each tree at each timepoint (each sampling point includes one enrichment and one direct plating sample)

|  | 21 March 2017 |  | 7 April 2017 |  | 12 June 2017 |  | 10 July 2017 |  |
| --- | --- | --- | --- | --- | --- | --- | --- | --- |
|  | soil | litter | soil | litter | soil | litter | soil | litter |
| Tree 1 | 4 | 4 | 2 | 2 | 4 | 4 | 4 | 4 |
| Tree 2 | 4 | 4 | 2 | 2 | 4 | 4 | 4 | 4 |
| Tree 3 | 4 | 4 | 2 | 2 | 4 | 4 | 4 | 4 |
| Tree 4 | 4 | 4 | 2 | 2 | 4 | 4 | 4 | 4 |
| Tree 5 | 4 | 4 |  |  |  |  | 4 | 4 |
| Tree 6 |  |  | 2 | 2 | 4 | 4 | 4 | 4 |
| Tree 7 |  |  | 2 | 2 | 4 | 4 | 4 | 4 |
| Tree 8 |  |  | 2 | 2 | 4 | 4 | 4 | 4 |
| Tree 9 |  |  |  |  | 4 | 4 | 4 | 4 |
| Tree 10 |  |  |  |  |  |  | 4 | 4 |

Material was collected simultaneously for the **direct plating** and **enrichment** collections at each sampling point (Figure 1). First, leaf litter was collected by aseptically transferring litter into sterile collection tubes: approximately 5 ml of compressed leaf litter was collected for the direct plating method and approximately 2 ml for the enrichment method. Then, the remaining leaf litter was removed from the soil surface and the top approximate 2 cm of soil (mostly composed of soil organic layer) were aseptically transferred into sterile collection tubes. As for leaf litter, approximately 5 ml of compressed soil was collected for the direct plating method and approximately 2 ml for the enrichment method. Instruments were sterilized between samples using 70% ethanol. Samples were transported between the field and lab at ambient temperature and processed within four hours of collection.

For **direct plating** (Figure 1A), material was mixed with 20 ml sterile water in a sterile 50 ml tube, the mixture was vigorously mixed for at least 10 seconds with a vortex mixer on its highest setting, and 0.2 ml of the resulting dirty liquid was pipetted on each of two plates containing the solid modified selective medium PIM1 (3 g yeast extract, 5 g peptone, 10 g sucrose, 3 g malt extract, 1 mg chloramphenicol, 80 ml ethanol, 5.2 ml 1 M HCl, and 20 g agar per liter) (Kowallik & Greig, 2016; Sniegowski et al., 2002). Liquid was spread on plates using sterile glass beads, and plates were left open in a laminar flow hood until dry. Plates were incubated for three days at 30 °C before colonies were picked.

For **enrichments** (Figure 1B), material was mixed with 10 ml of the liquid selective medium PIM1 (composition as for solid PIM1 but without agar) in a 15-ml sterile tube, mixtures were inverted, and tubes were incubated, slightly open and without shaking, at 30 °C. After 10 days, a sterile wooden stick was inserted into each enrichment tube and a small amount of the liquid (approximately 50 µl) was streaked onto a single plate containing the solid selective medium PIM2 (20 g Methyl-(alpha)- D-glucopyranoside, 1 ml 5% Antifoam Y-30 emulsion, 6.7 g Yeast Nitrogen Base without amino acids, 4 ml 1M HCl, and 20 g agar per liter) (Kowallik & Greig, 2016; Sniegowski et al., 2002), and plates were incubated 4 days at 30°C before colonies were picked.

We include these procedures as step-by-step protocols for the convenience of future researchers in the supplementary materials (Supplemental File 1).

### Yeast identification

After incubation, we streaked colonies with yeast-like morphology to fresh YPD medium (10 g yeast extract, 20 g peptone, 20 g dextrose, and 25 g agar per liter). For each method, up to 6 (March and April sampling days) or 12 (June and July sampling days) colonies per sample were selected. After one day of growth on YPD at 30 °C, cultures were frozen at −80 °C in 20% glycerol and a small amount of each culture was transferred to sporulation medium (20 g potassium acetate, 2.2g yeast extract, 0.5 g dextrose, 870 mg complete amino acid mixture, and 25 g agar per liter). Any cultures with bacteria-like morphology on YPD medium (slimy culture and/or cells smaller than 1 micron across) were not frozen and were discarded. Sporulation cultures were incubated for at least three days at room temperature before being screened under a compound microscope for *Saccharomyces*-like asci (tetrads).

All cultures producing tetrads were identified using sequencing of the internal transcribed sequence (ITS), a region neighboring rRNA-coding DNA (Schoch et al., 2012). We sequenced every strain using the ITS1/ITS4 primer pair (White, Bruns, Lee, & Taylor, 1990). PCR mixes were 7-15 µl in volume and contained one yeast colony, 0.5 µM each primer, and either 50% Phusion^®^ High-Fidelity PCR master mix with HF buffer or 1x HF-buffer, 100 µM dNTP mix, 3% DMSO, and 1 U/50 µl Phusion DNA polymerase. PCR reactions were cycled at 98 °C for 30 s, then 35 cycles of 98 °C for 5 s, 62 °C for 20 s, and 72 °C for 30 s, plus a 10 min terminal extension at 72 °C. PCR products were cleaned using illustra™ ExoProStar™ according to the manufacturer’s instructions, and sequenced on an ABI 3130xl sequencer.

ITS sequences were compared to sequences from the type or neotype strains of *S. paradoxus, S. cerevisiae, S. kudriavzevii*, and *S. mikatae* (Genbank accession numbers NR_138272.1, NR_111007.1, KY105195.1, and KY105198.1). If a sequence did not align with *Saccharomyces* sequences, we compared the sequence with all sequences in the NCBI database from type strains using BLAST (Zhang, Schwartz, Wagner, & Miller, 2000). If the sequence aligned with *Saccharomyces* sequences but had more than one base pair different from its closest match, we supplemented ITS sequences with sequences from the gene for translation elongation factor 1 using primers EF1-983F and EF1-2212R (Rehner & Buckley, 2005) using the protocols above, but with a PCR annealing temperature of 57 °C. In some cases, cultures originating from apparent single colonies were in fact mixtures of two yeast species. We counted these colonies as *Saccharomyces* if sequences from one of the species was *Saccharomyces*.

### Growth rates

We compared the distributions of maximum growth rates between the two groups of *S. paradoxus* strains (strains collected using enrichment culturing and strains collected using direct plating) in three liquid media. The media were liquid PIM1, a minimal yeast medium (1.7 g yeast nitrogen base without amino acids and ammonium, 5 g ammonium sulphate, and 2.5 g dextrose per liter), and liquid YPD (composition as for solid YPD, but without agar). To avoid confounding effects of environmental source (*i.e.*, combination of substrate, date collected, and tree), we compared growth rates for pairs of *S. paradoxus* strains originating from the same environmental source. In other words, we collected a dataset of *S. paradoxus* growth rates from two groups of strains with equal representations of combinations of substrate, date collected, and tree, and differing only in the method used to isolate the strains. To ensure that all isolates were pure *S. paradoxus* cultures (some cultures that came from what appeared to be single colonies during isolation were found to be mixtures of multiple species after ITS sequencing), we streaked all isolates used for growth rate measurements to single-colony cultures a second time. We confirmed that these single-colony cultures were *S. paradoxus* by mating them with a *S. paradoxus* tester strain (NCYC 3708, α, ura3::KANMX, ho::HYGMX). In total, 110 isolates (55 from each sampling method) were measured.

Growth rates were measured using an Epoch 2 microplate reader (Biotek Instrument, Inc., Winooski, VT, USA) and calculated using the included Gen5 software version 3.03.14 (Biotek Instrument, Inc., Winooski, VT, USA). We first inoculated strains in 0.2 ml of each liquid medium in a 96-well microplate and incubated cultures without shaking or measurement in the microplate reader at 30 °C for 24 hours to condition strains to microplate reader conditions. We then transferred 2 µl from each culture to 198 µl fresh medium in a new microplate and incubated the new microplate under the same conditions for 20-24 hours. OD_660_ was measured during the second incubation every ten minutes, and maximum growth rate (mOD_660_/min) was calculated from the maximum slope of each growth curve over four points (30 min total) using Gen5 software. Reported growth rates for each isolate are means of three replicates.

### Genotyping

Nine microsatellite loci were identified by searching for common *S. cerevisiae* repeats in the reference genome of *S. paradoxus* strain CBS432 (Young, Sloan, & Van Riper, 2000; Liti et al., 2009) and by adapting previously published *S. cerevisiae* microsatellite loci for *S. paradoxus* (González Techera, Jubany, Carrau, & Gaggero, 2001; Legras, Ruh, Merdinoglu, & Karst, 2005). Primers were designed using Primer3 2.3.4 in Geneious 8.1.8 (Untergasser et al., 2012, https://www.geneious.com). Seven microsatellite loci were three-nucleotide repeats; one locus was two-nucleotide repeats; and one locus was four-nucleotide repeats. All loci are described in Table 2. Some loci were complex, including repeats with different sequences; when analyzing data, we assumed that alleles of these loci with the same length had the same sequence.

**Table 2:**
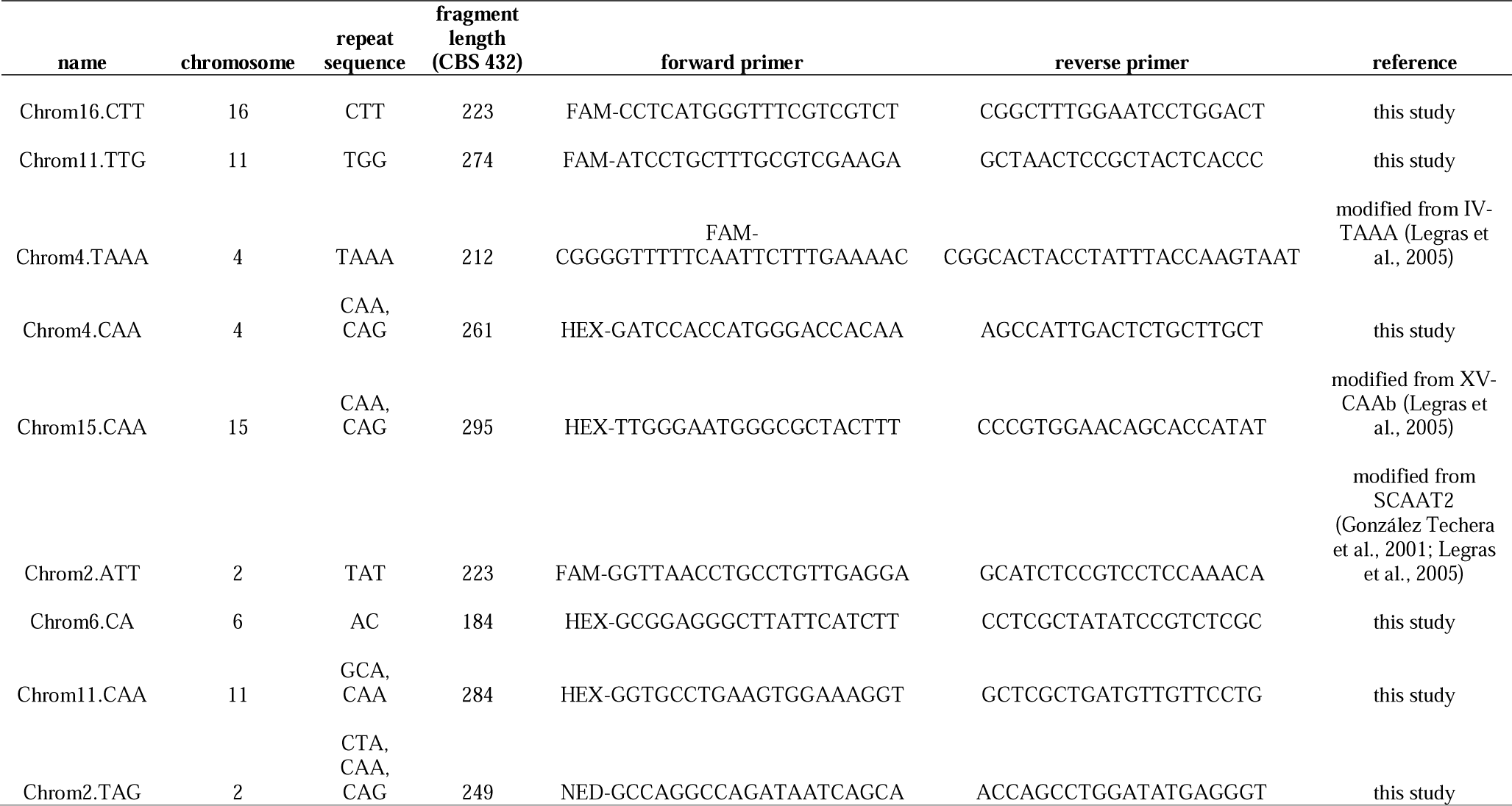
Information about microsatellite loci used to genotype *S. paradoxus* isolates

All *S. paradoxus* strains for which growth rates were measured (see above) were genotyped. We amplified microsatellite regions as previously described (Babiker & Tautz, 2015; Hardouin et al., 2015), with slight modifications. Reactions were carried out in 5 µl PCR mixes containing one colony of each *S. paradoxus* isolate, 2.5 µl 2x Qiagen Multiplex PCR master mix and 0.2 µM each primer. Forward primers were labeled with either FAM, HEX, or NED at the 5-prime end, and we multiplexed 4-5 primer pairs in each reaction. PCR cycling, dilution, and denaturation were carried out as previously described (Babiker & Tautz, 2015; Hardouin et al., 2015); fragments were run on an ABI 3730 DNA analyzer and were analyzed using Geneious 8.1.8 with microsatellite plugin version 1.4.4. Genescan ROX-500 (ThermoFisher Scientific) was used as a size standard. All nine microsatellite loci showed variation in the collection of *S. paradoxus* isolates: the lowest number of length polymorphisms detected for any locus was two and the maximum was ten.

### Statistical analyses

We compared sampling success across substrates (leaf litter or soil) and methods (direct plating or enrichment) using a generalized linear mixed-effects model with probability of isolating *Saccharomyces* (including both *S. paradoxus* and *S. cerevisiae*) as the response variable, substrate and method as fixed effects, and tree and date as random effects. We selected the best model using a top-down strategy, comparing Akaike’s Information Criteria (AIC) after removing predictors from a full model one by one. Models were calculated using the lme4 package in R version 3.6.0 (Bates, Machler, Bolker, & Walker, 2015; R Development Core Team, 2019).

We compared growth rate distributions by first comparing variances using Levene’s test for homogeneity of variance (Levene, 1960), and then comparing medians using paired Wilcoxon signed rank tests. We visualized relationships among genotypes using a neighbor-joining tree of Edwards genetic distance (Edwards, 1971). Genotypes detected per sample (excluding samples in which only a single isolate was measured) were compared between methods using a paired Wilcoxon signed rank test, and total genotypes isolated were compared between methods using the bootstrap method described in (Chao, et al. 2014) with 50 replications. Statistics were calculated using R version 3.6.0 (R Development Core Team, 2019) and the poppr, ape, car, and iNEXT packages (Fox, & Weisburg, 2019; Hsieh, Ma, & Chao, 2019; Kamvar, Tabima, & Grünwald, 2014; Paradis, & Schliep, 2018). Graphics were produced using the ggplot2 package and FigTree v.1.4.3 (Wickham, 2016, http://tree.bio.ed.ac.uk/software/figtree/).

## Results

### Influence of sampling method on *Saccharomyces* isolation success

Direct plating was more successful than enrichment culturing for isolating *Saccharomyces* spp. from natural substrates (z = 6.1, p < .001) (Tables 3, 4, Figure 2). We found *Saccharomyces* isolates in 45% of direct plating samples and 19% of enrichment culturing samples. However, enrichment culturing produced the only *S. cerevisiae* found in this study: we found six *S. cerevisiae* isolates from a single enrichment culture from tree 3 in March of 2017. All other *Saccharomyces* isolates found in this study were *S. paradoxus*. Other detected yeast species included *Saccharomycodes ludwigii, Torulaspora delbrueckii, Pichia membranifaciens*, and *Hanseniaspora osmophila*, all of which have previously been found alongside *Saccharomyces* spp. in beverage fermentations (Domizio et al., 2011; Gschaedler, 2017).

**Table 3:**
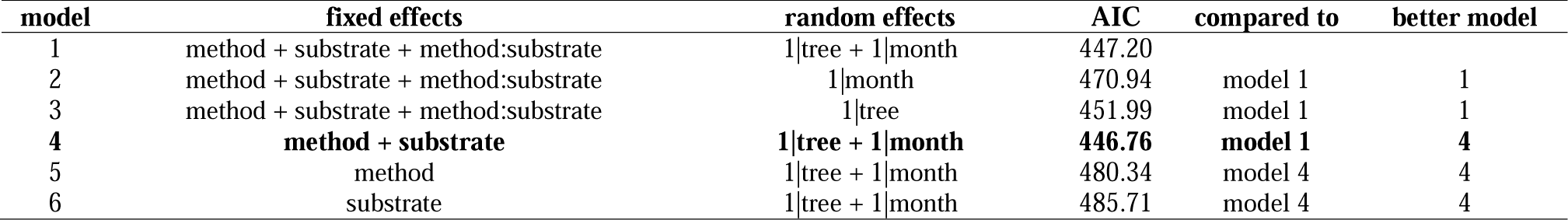
Model selection (Mixed-effects generalized linear model)

**Table 4:** Model summary table (model 4)

|  | Estimate | Std. Error | z | p |
| --- | --- | --- | --- | --- |
| intercept | -2.5324 | 0.4599 | -5.506 | < .001 |
| method (plating) | 1.5494 | 0.2559 | 6.054 | < .001 |
| substrate (soil) | 1.4445 | 0.2542 | 5.683 | < .001 |

**Figure 2:**
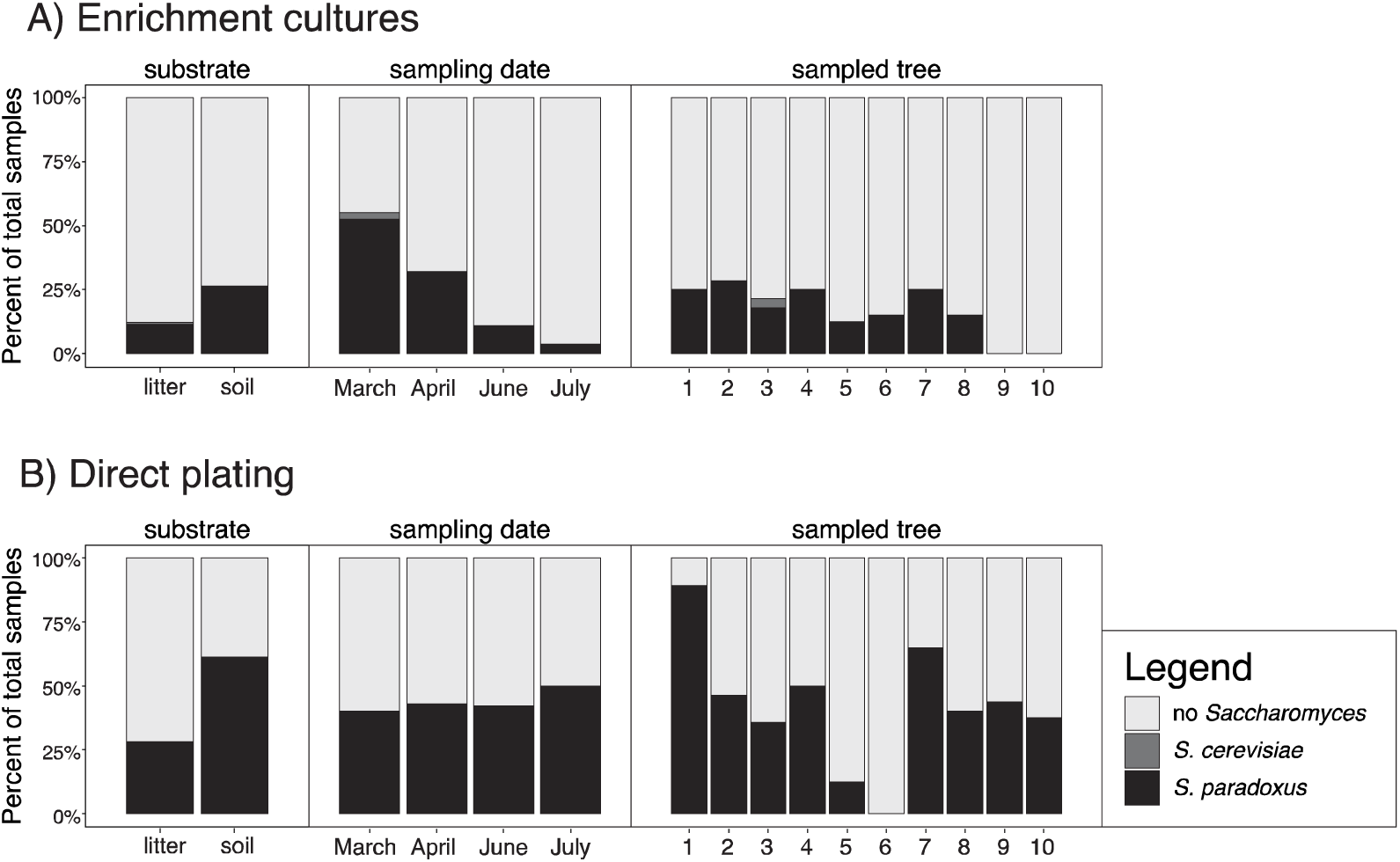
Percentages of samples in which *Saccharomyces* could be detected using A) enrichment cultures or B) direct plating. Bars represent all samples for each category of sampling, and shading represents *Saccharomyces* species. *S. paradoxus* and *S. cerevisiae* were the only detected *Saccharomyces* species.

While the direct plating method was more successful than the enrichment method, it was also more labor-intensive (Table 5). We found more colonies with *S. paradoxus*-like morphology, including colonies that belonged to non-*Saccharomyces* genera, using the direct plating method (969) than using the enrichment method (284), and we screened all of these colonies for tetrad formation. As a result, we screened more than three times as many colonies for tetrads when using the direct plating method than we did using the enrichment method. After screening for tetrads and ITS sequencing, only 32% of the total isolated direct plating colonies were *S. paradoxus*, compared to 74% of enrichment colonies.

**Table 5:**
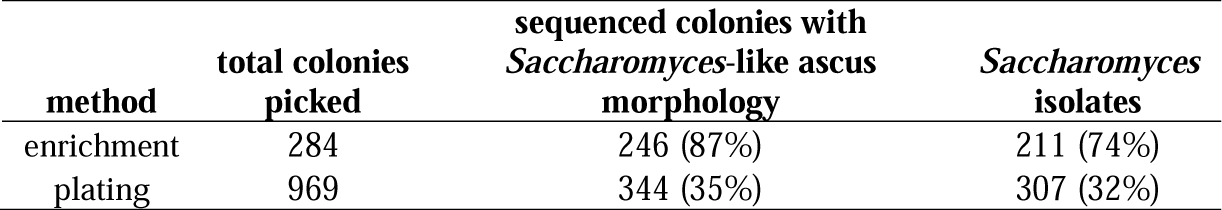
Colonies processed and sampling success for each method

Both methods isolated *Saccharomyces* colonies from both substrates, most trees, and all timepoints (Figure 2). We had significantly more sampling success on soil than leaf litter substrates (z = 5.7, p < .001, Table 4), but other relationships among sampling success, sampling method, and sampling environments were idiosyncratic. For example, direct plating did not produce any *Saccharomyces* isolates from tree 6, while three enrichment samples from this tree isolated *S. paradoxus*, and enrichments produced more *Saccharomyces* isolates in March than direct plating did (Figure 2). Because our sampling effort was not the same for all trees at all months, we did not model tree habitat or sampling month as fixed effects; instead, we modeled these parameters as random effects, and found that models including tree and month fit the data better than models without tree and month (Table 3).

### Phenotype diversity of sampled *S. paradoxus*

Growth rate distributions did not differ between the two methods in PIM1 and the minimal medium, and differed slightly in variance in the YPD medium (Figure 3, Tables 6-7). Median growth rates did not differ significantly between the two methods in any of the three tested media (Table 6), and variances in growth rate only differed significantly in YPD (Levene’s test F_1,108_ = 5.42, p = .022, Table 7), with enrichment cultures isolating a wider variance of *S. paradoxus* growth rates in YPD than direct plating (Figure 3C). When two outlier strains were removed (Figure 3C), this difference disappeared (F_1,106_ = 3.59, p =.06).

**Table 6:** Median growth rate comparisons between sampling methods

| tested medium | Wilcoxon signed rank test paired V | p |
| --- | --- | --- |
| PIM1 | 818.5 | .69 |
| minimal | 911.5 | .24 |
| YPD | 733 | .76 |

**Table 7:** Variance in growth rate comparisons between sampling methods

| tested medium | F | df | p |
| --- | --- | --- | --- |
| PIM1 | 0.99 | 1, 108 | .32 |
| minimal | 3.17 | 1, 108 | .078 |
| YPD | 5.42 | 1, 108 | .022 |

**Figure 3:**
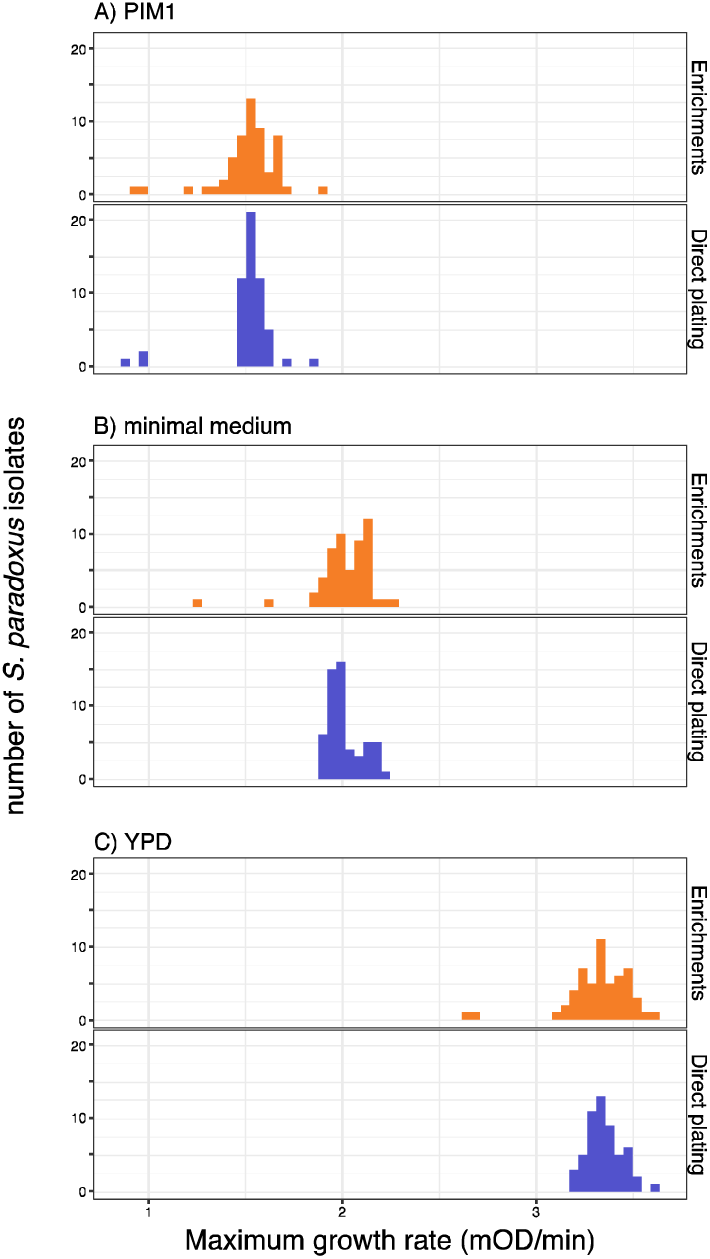
Histograms representing distributions of growth rates for *S. paradoxus* clones isolated using enrichment culturing and direct plating in A) PIM1, B) minimal, and C) YPD media.

### Genotype diversity of sampled *S. paradoxus*

The two isolation methods sampled equivalent genotype diversities, both across and within samples. In total, we found 21 unique clonal genotypes (Figure 4). The minimum number of clones per genotype was one and the maximum was 55 (Supplemental Figure 1). The enrichment method discovered 17 genotypes (95% confidence interval 12.1-21.9), and the direct plating method discovered 12 genotypes (95% confidence interval 8.8-15.2), but this difference was not significant because the two confidence intervals overlap (Figure 4). Enrichment cultures sampled more genotypes per sample (mean = 1.71, samples with only one isolate excluded) than direct plating cultures (mean = 1.57), but this difference was also not significant (Wilcoxon signed rank test paired V = 27, p = .61). All genotypes except one were homozygous at all loci. The single heterozygous genotype was present in two isolates sampled from leaf litter beneath tree 7 in June of 2017; the enrichment sampling and direct plating methods each isolated one of the two heterozygous isolates.

**Figure 4:**
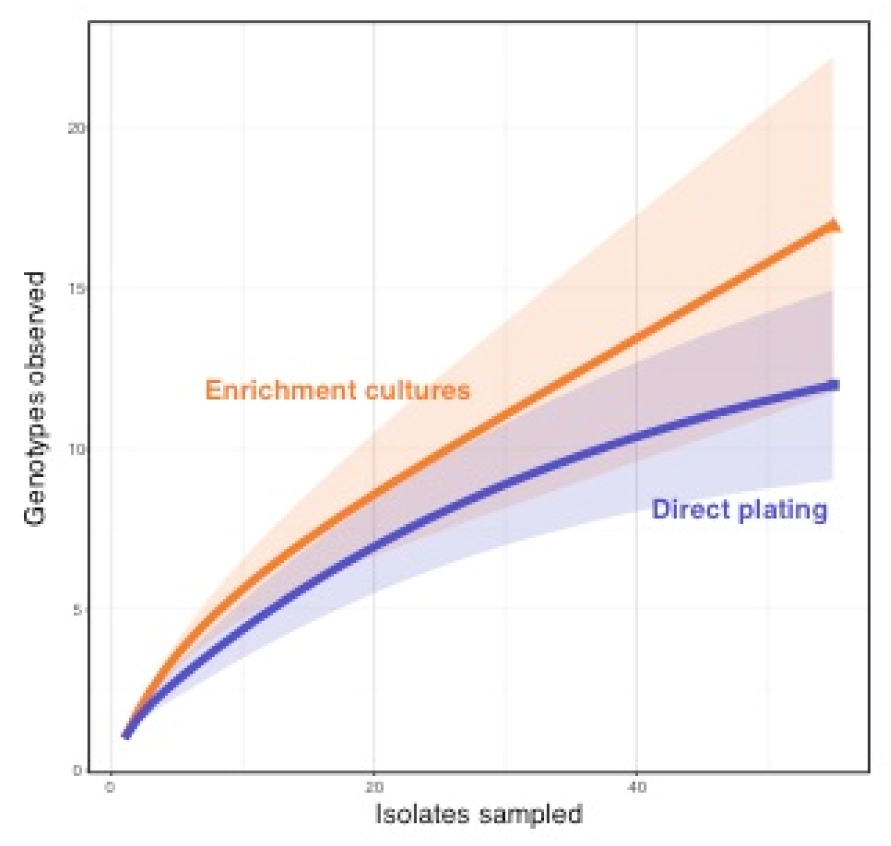
Genotype rarefaction curves of genotypes detected. Thick lines represent average genotypes observed as a function of isolates sampled and shated areas represent 95% confidence intervals.

## Discussion

### Direct plating detects *S. paradoxus* more frequently than enrichment culturing

Enrichment culturing did not increase *Saccharomyces* sampling success per collected forest leaf litter and soil sample compared to direct plating, in spite of researchers’ long history of using enrichment culturing to isolate *Saccharomyces* from forest environments (Kowallik & Greig, 2016; Naumov, Naumova, & Sniegowski, 1998; Sniegowski et al., 2002). We expect reliable *Saccharomyces* isolation from this forest using direct plating to be a result of high *S. paradoxus* abundance on forest floor substrates. Indeed, a previous study determined that hundreds to tens of thousands of *S. paradoxus* cells can occupy a gram of leaf litter near the bases of oak trees in this forest (Kowallik & Greig, 2016). These quantitative observations were made by serially diluting enrichment cultures and estimating the number of *S. paradoxus* cells per gram of leaf litter based on the highest dilution in which *S. paradoxus* could be found. We expect direct plating to be less successful in environments in which *Saccharomyces* are rarer, and note that enrichment culturing is frequently used to isolate *Saccharomyces* from tree bark, which may be a habitat with lower *Saccharomyces* density than the forest floor habitats we sampled (Kowallik et al., 2015; Sniegowski et al., 2002). *S. paradoxus* abundance can also vary over time, with spikes after environmental changes such as rain events (Anderson et al., 2018; Glushakova et al., 2007). It is possible that environmental conditions at other locations, or characteristics of non-European *S. paradoxus* populations, would result in different sampling successes using these two methods from that reported here.

It was not possible to completely standardize quantities of sampled natural material when comparing direct and enrichment-based sampling methods. We collected a larger volume of material for direct cultures (∼5 ml) than for enrichment cultures (∼2 ml), but the proportion of the original enrichment sample ultimately screened for *Saccharomyces* colonies depends on processes occurring during enrichment. For direct plating, we screened 400 µl of the 25 ml total suspension of soil or leaf litter material in water for *Saccharomyces* colonies. For enrichment culturing, we potentially screened all 2 ml of collected material (if *S. paradoxus* cells present in low cell numbers at the start of enrichment culturing grew to high cell numbers during the enrichment incubation) or none of the collected material (if other microbes in enrichments inhibited *S. paradoxus* growth). *S. paradoxus* sampling success in enrichment cultures depends on the composition of the co-sampled microbial community, and as a result we chose to standardize by maximum number of colonies screened (6 or 12) instead of by volume of material collected. Researchers adapting our methods could adjust the amount of material collected, the volume of liquid plated, or the number of colonies screened to optimize the methods to their own systems.

### Both isolation methods sampled similar *S. paradoxus* diversities from forest substrates

Overall, enrichment culturing and direct plating collected similar phenotypic and genotypic diversity (Figures 3, 4). There was also no evidence that enrichment culturing selected for individuals that were fitter under enrichment conditions than individuals sampled using direct plating (Figure 3A). While we genotyped many representatives of the same clonal genotypes (Figure S1), clonal reproduction inside of enrichment cultures did not decrease sampled diversity. High clonality in a local area is common for wild *S. paradoxus* populations, and is most likely a result of extensive asexual reproduction in natural habitats (Tsai, Bensasson, Burt, & Koufopanou, 2008; Xia, et al. 2017). We also found no evidence for sexual outcrossing in the enrichment cultures themselves. If outcrossing had occurred during enrichment, we would expect to have seen heterozygous F1 offspring among the genotypes isolated using enrichment. Instead, the only heterozygous genotype in our collection was isolated from a single environment using both enrichment and plating methods, and was unlikely to have arisen during enrichment.

The enrichment method did isolate some outlier *S. paradoxus* phenotypes and other *S. cerevisiae* individuals that the direct plating method did not (Figures 2, 3C). We did not find many of these outliers, but we speculate that diverse interactions with microbes in enrichments may have led to isolation of outlier phenotypes and *Saccharomyces* spp. For example, the isolated outlier *Saccharomyces* may have come from enrichments containing bacteria that promoted outlier *S. paradoxus* or *S. cerevisiae* growth at the expense of other *S. paradoxus* genotypes. Microbial diversity across enrichment cultures may similarly explain our idiosyncratic sampling success across months and trees (Figure 2). For example, it is possible that a bacterium that inhibits *S. paradoxus* growth in the enrichment medium was more common in summer than spring months, resulting in lower enrichment sampling success in summer.

### Recommendations for future yeast sampling

Our results identified a tradeoff between resources spent on sampling and resources spent on sequencing: enrichment culturing was less successful than direct plating at finding *Saccharomyces* per sample collected, but more successful per ITS region sequenced (Figure 2, Table 5). Researchers with a few precious samples are therefore better off isolating *Saccharomyces* using direct plating, especially if *Saccharomyces* is common on their substrates. Conversely, if samples are easy to get but funds available for sequencing are limited, researchers may prefer to use enrichment culturing or to use direct plating with more phenotypic screening tests than we used. For example, researchers using the direct plating protocol might replica-plate colonies to a second selective medium such as PIM2, which did not increase sampling biases in the enrichment cultures in our sampling, to reduce the number of non-*Saccharomyces* colonies that must be sequenced. Researchers with limited time or freezer space who would like assurances that most picked colonies are *Saccharomyces* may also prefer enrichment culturing or direct plating with additional selective steps.

While, on average, both methods sampled similar phenotypic and genotypic diversity, our isolation of outlier isolates using enrichments suggests that researchers targeting outliers may also prefer enrichment culturing. For example, researchers sampling environments to find unusual *Saccharomyces* phenotypes for applied biotechnology (*e.g.*, food microbiology, drug discovery) may uncover more diversity using enrichment culturing. Researchers interested in detecting rare *Saccharomyces* species in an environment (*e.g., S. cerevisiae* from our study forest, *S. mikatae* and *S. eubayanus* from European forests) (Alsammar et al., 2018) may also have more success using enrichment culturing.

## Conclusions

Our results validated use of enrichment culturing for isolating diverse and representative collections of *S. paradoxus* from natural material. We found no evidence that processes during enrichment culturing decrease the diversity of sampled *Saccharomyces* spp., and weak evidence that these processes may in fact increase sampled diversity. While it is generally a good idea to standardize sampling methods within a study as much as possible, conclusions from studies comparing *Saccharomyces* genotype and phenotype diversity from a variety of sources, including culture collections, are likely to be reliable (Strope et al., 2015; Warringer et al., 2011) and the diversity found in culture collections is likely to be representative of natural *Saccharomyces* diversity in sampled environments. In addition to validating the frequently-used enrichment method for isolating *Saccharomyces* spp., this study provides a reliable direct method for isolating *Saccharomyces* spp. and describes a set of microsatellite markers that can be used to conveniently identify *S. paradoxus* genotype diversity. The utility of *Saccharomyces* as an ecology and evolutionary model relies on our understanding of its natural history, and we hope that these and other improvements in field sampling methods will empower researchers the explore the environmental contexts of these exciting microbial model organisms.

## Acknowledgements

We would like to thank Danielle Stevens and Tjorben Nawroth for help in the field, Jenna Gallie and the Gallie lab for help with growth curves, Michael Habig for advice on statistics, and Amine Hassani for helpful conversations on microbial diversity in enrichment cultures. Thank you to Christoph Freiherr von Fürstenberg-Plessen for permission to work in the Nehmten forest. This work was supported through a Max Planck Fellowship to Eva H. Stukenbrock.

## Data Accessibility

All data for this project, including a list of identified yeasts, sampling success data for each soil and leaf litter sample, isolate growth rates, and isolate genotype data have been deposited in the Edmond repository (doi:10.17617/3.2i).

## Figure Legends

Supplemental Figure 1: Neighbor-joining tree of isolates genotyped for this study. The scale bar reflects Edwards distance (9 loci scored). The tree was rooted with N-44 (CBS 8438, a S. paradoxus strain from East Asia). The red circle indicates two heterozygous isolates (heterozygous at two of the nine loci); all other isolates were homozygous at all loci.

