## Supplemental Figure 1 for "Quantifying the efficiency and biases of forest *Saccharomyces* sampling strategies"

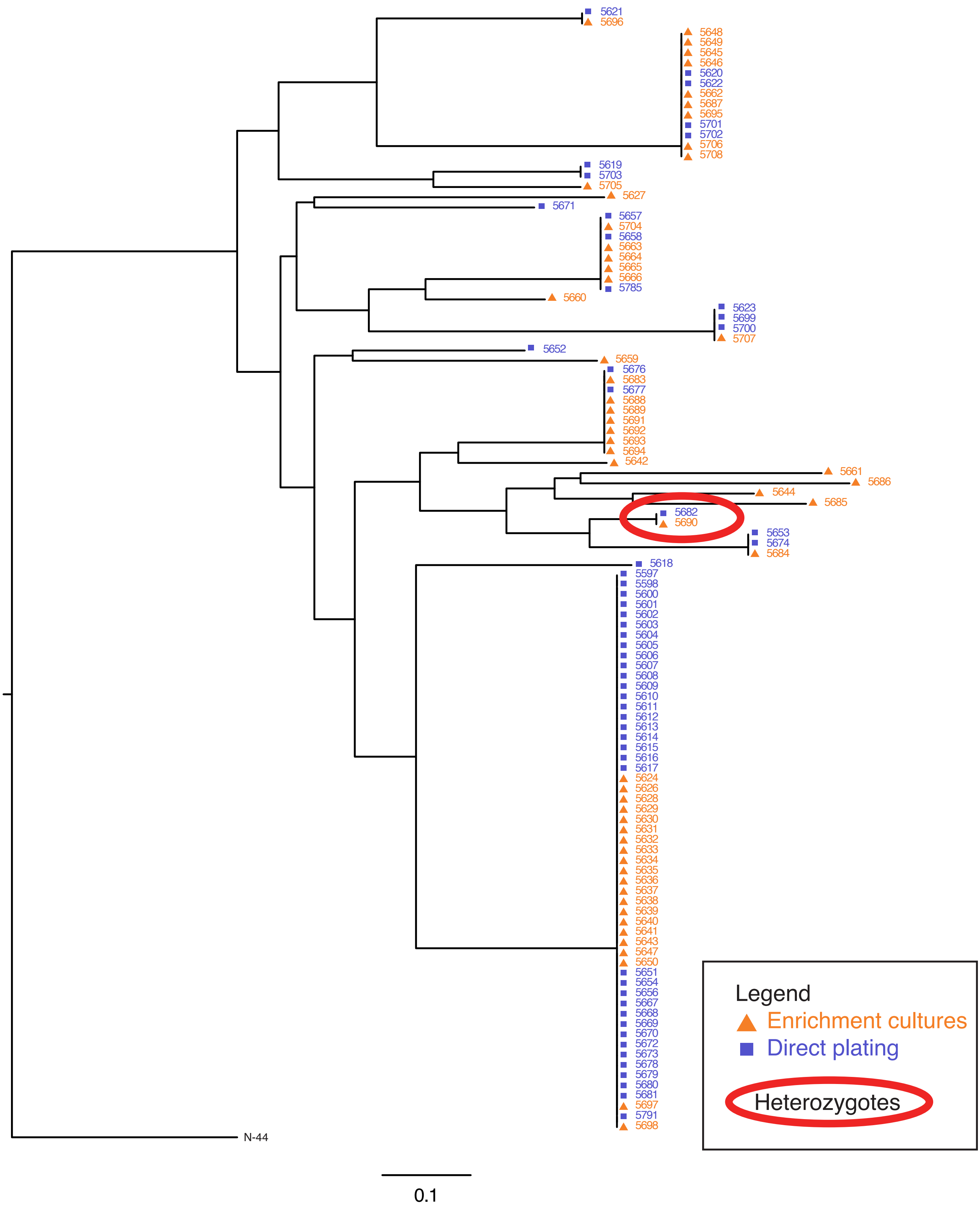


Suplemental Figure 1: Neighbor-joining tree of isolates genotyped for this study. The scale bar reflects Edwards distance (9 loci scored). The tree was rooted with N-44 (CBS 8438, a *S. paradoxus* strain from East Asia). The red circle indicates two heterozygous isolates (heterozygous at two of the nine loci); all other isolates were homozygous at all loci.
