## Supplemental File 1 for "Quantifying the efficiency and biases of forest *Saccharomyces* sampling strategies"

Protocols for sampling *Saccharomyces* from forest leaf litter and soil

**Enrichment culturing protocol**

(adapted from Sniegowski *et al.* 2002 and Kowallik & Greig 2016)

Materials needed (media recipes below)

- sterile 15-ml tubes (1 per sample, labeled in advance if possible)
- Sterile plastic containers for holding spatulas and forceps between sampling (large pipette tip boxes work well)
- At least 2 spatulas or spoons, autoclaved (choose tools that can fit into the 15-ml tubes)
- At least 2 large forceps (~15 cm long with blunt tips), autoclaved
- Several 50-ml tubes half full of 70% ethanol
- Liquid PIM1 media (10 ml per sample)
- Pipette with at least 10-ml capacity and bulb or pump
- PIM2 plates (1 per sample)
- Sterile long wooden sticks (long enough to fit in the 15-ml tubes) or other tool for streaking samples
- YEPD plates
- Sampling notebook

Protocol

*Field sampling steps*

1. For each leaf litter sample, open one 15-ml tube and place mouth-down on the leaf litter surface. Do this as close as possible to the tree trunk, and not more than 1 m from the base of the tree
2. Use the tube and sterilized forceps to push about 2 cubic centimeters of compressed leaf litter into the tube. Try to sample entire “litter horizon”, *i.e.*, all litter between the litter surface and soil surface
3. Use the forceps to push remaining leaf litter from the top of the soil surface, exposing a square of soil about 5 x 5 cm
4. For each soil sample, open another 15-ml tube and place mouth-down on the soil surface. Press down about 2 cm into the top layer of soil.
5. Use the tube and sterilized spatula to push about 2 cubic centimeters of compressed soil into the tube.
6. Close all sampling tube lids
7. Sterilize tools between samples by agitating them in 70% ethanol, and let them dry by resting them in sterile plastic containers
8. Record the tree, substrate, and other information (*e.g.*, compass direction relative to tree trunk, distance from tree trunk)
9. Transport tubes at ambient temperature and process within four hours

*Lab culturing steps*

1. Check to make sure ethanol, antibiotics, and acid have been added to PIM1 (it is better to add these ingredients to the media immediately before use)
2. Add 10 ml of liquid PIM1 to each 15-ml sampling tube
3. Invert to mix, exposing all sampled material to the media, and tap each tube to remove air bubbles from the bottom
4. Place lids on sample tubes, but do not tighten lids. Allow gasses to escape
5. Incubate without shaking at 30 ºC
6. After 10 days, dip a sterile wooden stick (or other sterile instrument) into each tube and streak out onto a PIM2 plate. Incubate PIM2 plates 4 days at 30 ºC
7. Streak up to 6 or up to 12 colonies to YEPD plates and grow overnight at 30 ºC
8. Freeze cultures for storage (-70 to -80 ºC) and identify morphologically or using DNA sequences

**Direct plating protocol**

(adapted from above protocol and Kowallik 2015)

Materials needed

- sterile 50-ml tubes (1 per sample, labeled in advance if possible)
- at least 1 L sterile water (20 ml per sample)
- Sterile 0.5-cm glass beads
- pipette and sterile pipette tips (1 ml capacity is best)
- Sterile plastic containers for holding spatulas and forceps (large pipette tip boxes work well)
- At least 2 spatulas or spoons, autoclaved (choose tools that can fit into the 50-ml tubes)
- At least 2 large forceps (~15 cm long with blunt tips), autoclaved
- Several 50-ml tubes half full of 70% ethanol
- Vortex mixer
- Pipette with at least 20-ml capacity and bulb or pump
- PIM1 plates with 2% agar (2 per sample)
- YEPD plates
- Sampling notebook

Protocol

*Field sampling steps*

1. For each leaf litter sample, open one 50-ml tube and place mouth-down on the leaf litter surface. Do this as close as possible to the tree trunk, and not more than 1 m from the base of the tree
2. Use the tube and sterilized forceps to push about 5 cubic centimeters of compressed leaf litter into the tube. Try to sample entire “litter horizon”, *i.e.*, all litter between the litter surface and soil surface
3. Use the forceps to push remaining leaf litter from the top of the soil surface, exposing a square of soil about 5 x 5 cm
4. For each soil sample, open another 50-ml tube and place mouth-down on the soil surface. Press down about 2 cm into the top layer of soil.
5. Use the tube and sterilized spatula to push about 5 cubic centimeters of compressed soil into the tube.
6. Close all sampling tube lids
7. Sterilize tools between samples by agitating them in 70% ethanol, and let them dry by resting them in sterile plastic containers
8. Record the tree, substrate, and other information (*e.g.*, compass direction relative to tree trunk, distance from tree trunk)
9. Transport tubes at ambient temperature and process within four hours

*Lab culturing steps*

1. Add 20 ml sterile water to each tube. Close lids tightly
2. Vortex as thoroughly as possible. Use the highest setting of the vortex mixer and vortex at least 10 seconds. Turn the tube during mixing so that its bottom and sides are in contact with the mixer
3. Pipette 200µl of the dirty water to each of two PIM1 plates holding 6-8 sterile glass beads.
4. Shake each plate so that the beads roll across the agar surface and spread the liquid evenly
5. Dry the plates in a sterile containment hood right side up if necessary
6. Turn plates upside-down, remove the beads, and incubate at 30 ºC
7. After 3 days, streak up to 6 or 12 colonies per sample with yeast-like morphology to YEPD, and grow overnight at 30 ºC
8. Freeze cultures for storage (-70 to -80 ºC) and identify morphologically or using DNA sequences

**Media recipes**

liquid PIM1 (Sniegowski *et al.* 2002, Kowallik & Greig 2016)

1. Mix 3 g yeast extract, 5 g peptone, 10 g sucrose, and 3 g malt extract with deionized water for a total volume of 1 L
2. Autoclave
3. Just before use, add 54 µl chloramphenicol solution (20 mg/ml in ethanol), 80 ml 96% ethanol, and 2.6 ml 2 M HCl, and mix

solid PIM1

1. Mix 3 g yeast extract, 5 g peptone, 10 g sucrose, 3 g malt extract, and 20 g agar with deionized water for a total volume of 1 L
2. Autoclave
3. Cool on a stir plate while gently stirring at room temperature until sides of the flask can be comfortably touched
4. Mix in 54 µl chloramphenicol solution (20 mg/ml in ethanol), 80 ml 96% ethanol, and 2.6 ml 2 M HCl. Add the ethanol slowly as it can cool the agar solution
5. Pour plates
6. Use plates within 1 week

PIM2 (Sniegowski *et al.* 2002, Kowallik & Greig 2016)

1. Mix 20 g Methyl-(alpha)-D-glucopyranoside, 6.7 g yeast nitrogen base without amino acids, 20 g agar, and 1 ml 5% Antifoam Y-30 with deionized water for a total volume of 1 L
2. Autoclave
3. Cool on a stir plate while gently stirring at room temperature until sides of the flask can be comfortably touched
4. Mix in 2 ml of 2 M HCl
5. Pour plates

YEPD

1. Mix 10 g yeast extract, 20 g peptone, 20 g dextrose, and 25 g agar with deionized water for a total volume of 1 L
2. Autoclave
3. Pour plates
